# Extracting and characterizing protein-free megabasepair DNA for *in vitro* experiments

**DOI:** 10.1101/2022.06.22.497140

**Authors:** Martin Holub, Anthony Birnie, Aleksandre Japaridze, Jaco van der Torre, Maxime den Ridder, Carol de Ram, Martin Pabst, Cees Dekker

## Abstract

Chromosome structure and function is studied in cells using imaging and chromosome-conformation-based methods as well as *in vitro* with a range of single-molecule techniques. Here we present a method to obtain genome-size (megabasepair length) deproteinated DNA for *in vitro* studies, which provides DNA substrates that are two orders of magnitude longer than typically studied in single-molecule experiments. We isolated chromosomes from bacterial cells and enzymatically digested the native proteins. Mass spectrometry indicated that 97-100% of DNA-binding proteins are removed from the sample. Upon protein removal, we observed an increase in the radius of gyration of the DNA polymers, while quantification of the fluorescence intensities showed that the length of the DNA objects remained megabasepair sized. In first proof-of-concept experiments using these deproteinated long DNA molecules, we observed DNA compaction upon adding the DNA-binding protein Fis or PEG crowding agents and showed that it is possible to track the motion of a fluorescently labelled DNA locus. These results indicate the practical feasibility of a ‘genome-in-a-box’ approach to study chromosome organization from the bottom up.

## Introduction

Over the past decade, bottom-up synthetic cell research or ‘bottom-up biology’ has gained traction as a method to study components of living systems. The ultimate aim of such efforts is to build a synthetic cell by assembling biological functionalities from the bottom up. This involves the reconstitution of the various parts of living cells from a set of well-characterized but lifeless molecules such as DNA and proteins.(Schwille, 2015) While the end goal of building a functional synthetic cell is yet far off, the bottom-up approach has already successfully been applied to constitute and study minimal cellular systems, for example, intracellular pattern formation (Litschel et al., 2018), cell division (Ganzinger et al., 2020), the cytoskeleton (Litschel et al., 2021), and cellular communication (Joesaar et al., 2019).

For studying chromosome organization in the eukaryotic nucleus or in bacterial cells, numerous studies have been made on live or fixed cells through imaging (Bintu et al., 2018; Ricci et al., 2015), chromosome conformation capture techniques (Brandão et al., 2021; Falk et al., 2019), *etc*., while *in vitro* protein-DNA interactions are often characterized at the single-molecule level using techniques such as Atomic Force Microscopy (Dame et al., 2000; Japaridze et al., 2017; Liang et al., 2017), magnetic (Kaczmarczyk et al., 2020; Sun et al., 2013) and optical tweezers (Lin et al., 2021; Renger et al., 2022), and DNA visualization assays (Davidson et al., 2019; Ganji et al., 2018; Golfier et al., 2020; Greene et al.; Kim et al., 2019). While these complementary approaches have yielded great insights, they leave a significant gap since typical single-molecule methods address the ~kilobasepair (kbp) scale while actual genomes consist of 10^5^ – 10^11^ bp long DNA. It would be useful to study genome-size DNA with bottom-up *in vitro* methods, including the emergent collective behavior associated with this length scale. We propose that such experiments, which we coin a ‘genome-in-a-box’ (GenBox) approach (Birnie and Dekker, 2021), may provide valuable insights into chromosome organization, somewhat analogous to the ‘particle-in-a-box’ experiments in physics which proved a useful abstraction to understand basic phenomena in quantum mechanics. However, such a GenBox method has so far been lacking due to an unavailability of methodologies to prepare long DNA substrates. Expanding from the kbp to the Mbp scale poses technical challenges, both in the handling of long DNA that is prone to shearing, and in the availability of long DNA, as common *in vitro* experiments are done on viral DNA (such as the 48.5 kbp lambda-phage DNA) which however is limited in length.

Here, we present a methodology for the extraction of chromosomal DNA from *E. coli* bacteria and the subsequent removal of native proteins, resulting in deproteinated DNA of megabasepair size which can be used for *in vitro* bottom-up experiments to study chromosome organization (Figure 1). We describe the extraction and purification protocol, characterize the DNA objects obtained, and present some first proof-of-principle experiments.

**Figure 1.**
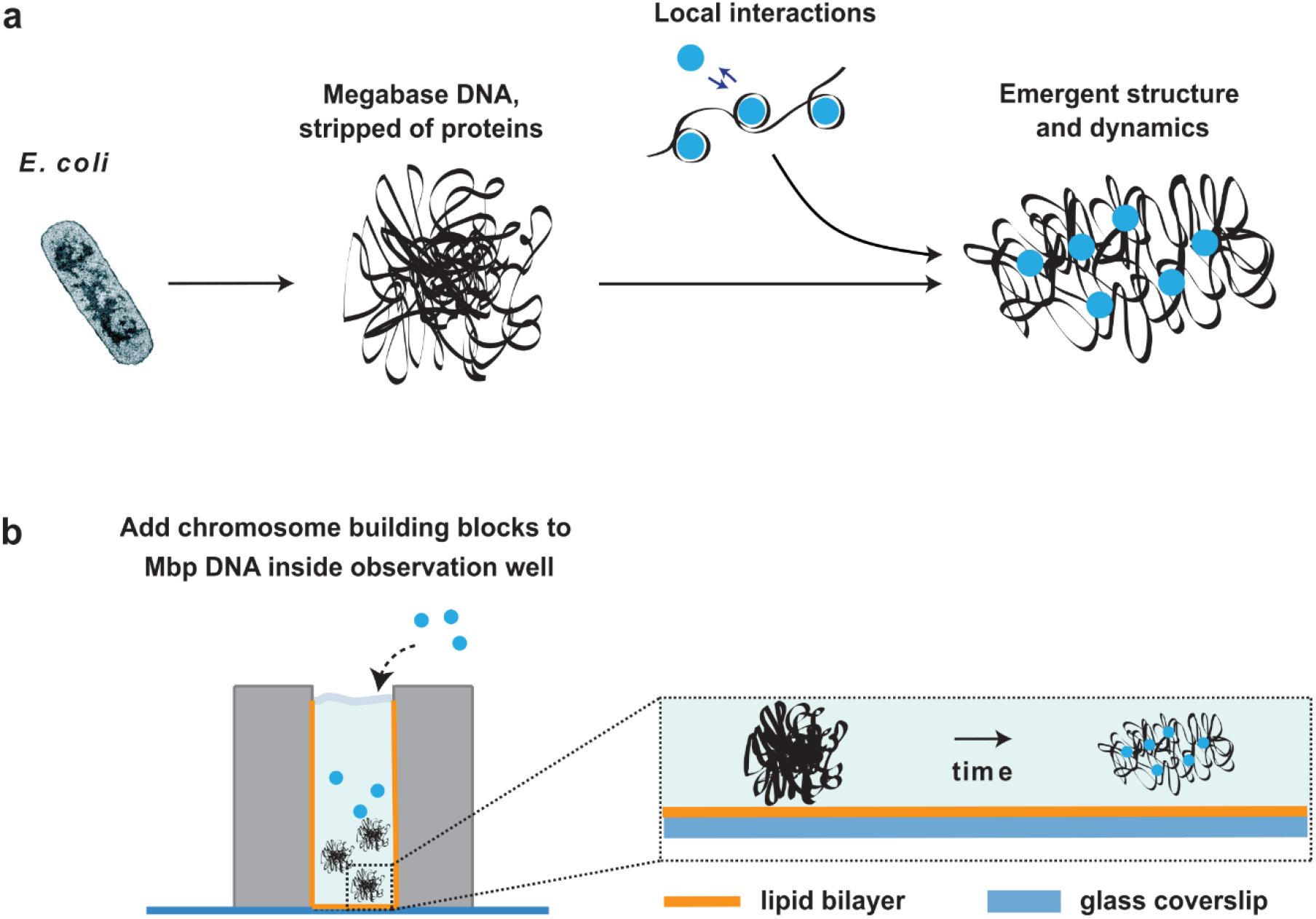
Methodology of extracting, purifying, and studying a bacterial chromosome. **a)** In a Genome in a box (GenBox) approach, one isolates chromosomes from bacterial cells, removes the natively bound proteins, to subsequently add DNA-structuring elements and thus study the resulting emergent DNA structure. **b)** Typical setup where a deproteinated megabasepair-long DNA is suspended in solution in an observation well attached to a glass coverslip. The surface of the observation well is coated with a lipid bilayer to prevent DNA adhesion to the surface. DNA-binding elements are added and the resulting DNA structure is observed using fluorescence microscopy.

## Results

The workflow to obtain and characterize deproteinated megabasepair DNA consisted of several experimental steps, which are discussed in the following sections. First, we ensured and verified that the *E. coli* bacteria contained a single 4.6 Mbp chromosome by cell-cycle arrest. Then chromosomes were extracted from the cells in one of two routes, either directly in solution or *via* embedding them in an agarose gel plug. Lastly, the isolated chromosomes were deproteinated using a protease enzyme. Mass spectrometry was used to confirm the level of deproteination, followed by microscopy imaging and quantitative analysis of the total fluorescence intensity per object and the radius of gyration (*R_g_*). This was done in order to verify if the chromosomes remained intact throughout the protocol, as well as to assess the effect of deproteination of the size of the DNA objects. Finally, as a proof of concept, three examples of possible experiments are shown.

### Extracting a single chromosome from *E. coli*

We prepared *E. coli* cells that contain only a single chromosome. In the exponential growth phase of bacteria, chromosomes are permanently replicating and typically exhibiting multiple replication forks on the DNA. For the purpose of controlled *in vitro* experiments this is undesirable for two reasons: first, halfway replicated DNA and multiple replication forks make the exact amount of DNA per cell unknown, and second, DNA near replication forks is prone to damage and breaking (Merrikh et al., 2012). As our aim is to extract DNA of a well-defined size, it is needed to obtain conditions that yield a known number of chromosomes per cell, ideally only a single chromosome per cell.

For this purpose, we used minimal media to avoid the occurrence of nested replication forks (Bird et al., 1972) as well as a temperature-sensitive *E. coli* strain where replication initiation was arrested by culturing the cells at an elevated temperature (Japaridze et al., 2020; Saifi and Ferat, 2012). We grew cells for 2 hours (*i.e*., for a time period longer than the doubling time in minimal media) at 41 °C and subsequently determined the number of chromosomes per cell by fluorescence imaging. The *E. coli* cells were engineered to contain Fluorescent Repressor Activator System (FROS) arrays near the Origin (Ori) and Terminus (Ter) locations (Figure 2a-*i*). At the start of the DNA replication process, the Ori is duplicated upon which the remainder of the chromosome follows, while the Ter is only duplicated at the end. This means that cells with a partly replicated chromosome will contain two Ori spots and a single Ter spot, whereas cells containing a single chromosome will only show one Ori and Ter. By counting the Ori and Ter fluorescence spots per cell, we confirmed that 85% of cells contained a single chromosome (Figure 2a-*ii* and *iii*), while 15% of cells were still in the process of DNA replication. If one were to extract the DNA from these cells, one would therefore expect a size distribution in which 85% of the objects are 4.6 Mbp, whereas the remaining 15% would contain DNA at an amount of between 4.6 and 9.2 Mbp, depending on how far genome replication in the cell had proceeded at the time of DNA extraction.

**Figure 2.**
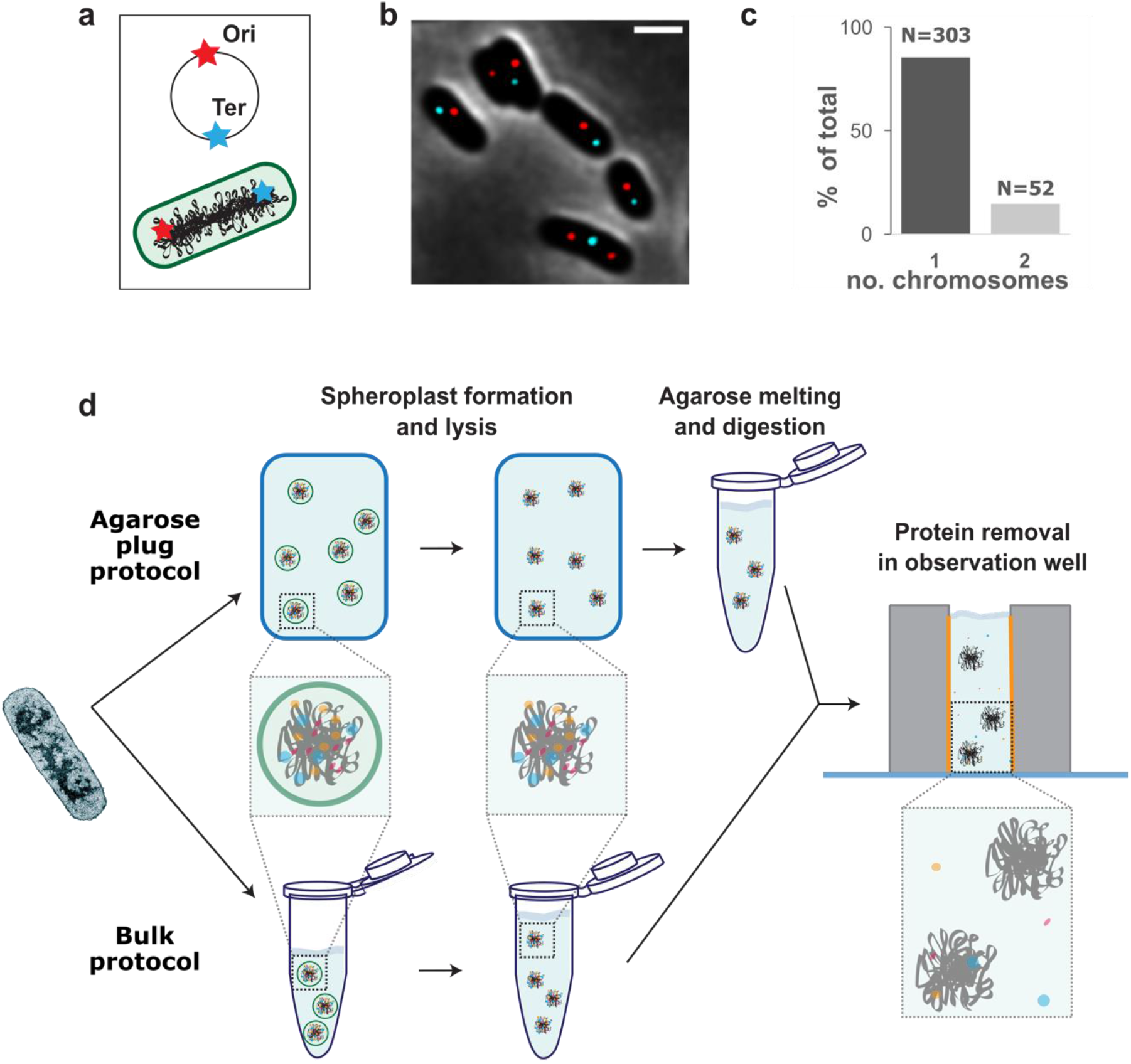
Workflow of the protocol. **a)** The *E. coli* chromosome is circular and contains FROS arrays near the Origin of replication (Ori) and Terminus of replication (Ter). **b)** Deconvolved image of *E. coli* cells with the Ori and Ter location labeled in red and cyan, respectively. Scale bar 2 μm. **c)** Number of chromosomes in temperature-treated *E. coli* cells. **d)** Agarose plug and bulk protocol to prepare deproteinated megabasepair DNA. Starting from *E. coli* cells, the cell wall is digested and the resulting spheroplasts are either embedded and lysed inside an agarose plug or directly lysed in a solution. After lysis in the agarose plugs, the agarose matrix is digested. At this stage, the chromosomes in both protocols are suspended in a solution and transferred to an observation well for protein removal and study of the deproteinated genomes.

In order to extract the chromosomes from *E. coli* cells, the peptidoglycan cell wall was degraded using lysozyme enzyme, resulting in spheroplasts which are wall-less rounded *E. coli* cells that merely are contained in their plasma membranes. To release the cellular contents including the DNA, the spheroplasts were submerged in a low-osmolarity buffer, which forces water to enter the spheroplasts, thereby rupturing them. This so-called lysis by osmotic shock was achieved on spheroplasts that were prepared with one of two methods (Figure 2b): *i*) direct lysis of the cytosolic content of the spheroplasts into solution, based on a protocol developed in the Woldringh lab (Cunha et al., 2001; Wegner et al., 2012) (hereafter called ‘bulk protocol’), or *ii*) embedding of spheroplasts inside agarose gel plugs where they were subsequently lysed, following a protocol from the Glass lab (Lartigue et al., 2007) (hereafter called ‘agarose plug protocol’). Embedding of the spheroplasts inside the agarose plug resulted in intact spheroplasts that did not get lysed prematurely (Figure S1). Bulk isolation yielded DNA that could be used on the same day, while the agarose-plug protocol produced samples that could be stored for a period of up to weeks after isolation. Depending on the application, the agarose plug protocol may also present advantages regarding the handing of the DNA material, such as a reduced shearing in transferring between experimental steps.

### Virtually all proteins can be removed from extracted chromosomes

DNA in cells is compacted by confinement, crowding, and binding of DNA-associated proteins. After cell lysis, the boundary conditions of confinement and crowding no longer apply, but DNA-binding proteins can in principle remain attached to the DNA. To digest such DNA-binding proteins in the sample, we incubated the bulk and plug protocol samples with a thermolabile Proteinase K enzyme, which is a broad range serine protease that cleaves peptide bonds at the carboxylic sides at a variety of positions (*i.e*., after aliphatic, aromatic, and hydrophobic amino acids). We observed increased DNA fragmentation after digesting and melting agarose plugs that had undergone proteinase treatment. Contrary to previous work (Lartigue et al., 2007), we therefore opted for treating the agarose sample in liquid, instead of in the gel state. While the bulk protocol sample already was liquid, agarose plugs had to be first digested using beta-agarase enzyme that breaks down the polymers forming the agarose gel. After the 15 min deproteination treatment and subsequent enzyme heat-inactivation (to prevent protein digestion in downstream experiments), we quantified the resulting degree of protein removal by mass spectrometry (MS).

Two categories of proteins were distinguished in the MS experiments, namely DNA-binding proteins and non-DNA-binding proteins. Obviously, the removal of the DNA-binding proteins is most critical for obtaining deproteinated DNA for GenBox experiments. To aid the quantification, we compiled a list of the 38 most abundant DNA-binding proteins as well as DNA-binding protein sub-units (Table S2), based on the protein’s description in the UniProt database as DNA-binding or DNA processing. For the bulk protocol (Table 1a-top), we found that all DNA-binding proteins were removed (100%, at the MS resolution). For the agarose plug protocol (Table 1b-top), the vast majority of the DNA-binding proteins, 97 %, was removed. These percentages refer to protein abundances relative to control samples that underwent exactly the same treatment steps, but to which no Proteinase K was added. For the agarose plug protocol (Table 1b-bottom), the major remaining DNA-binding proteins were IHF-A (14.8% remaining) and various RNA polymerase sub-units (rpoA/B/C, up to 4.5% remaining). The non-DNA-binding proteins were removed to the degree of 98.1% and 93.0% for the bulk and agarose plug protocol, respectively. More specifically, several ribosomal proteins were still present at large percentages (>40%) in the agarose plug sample.

**Table 1.**
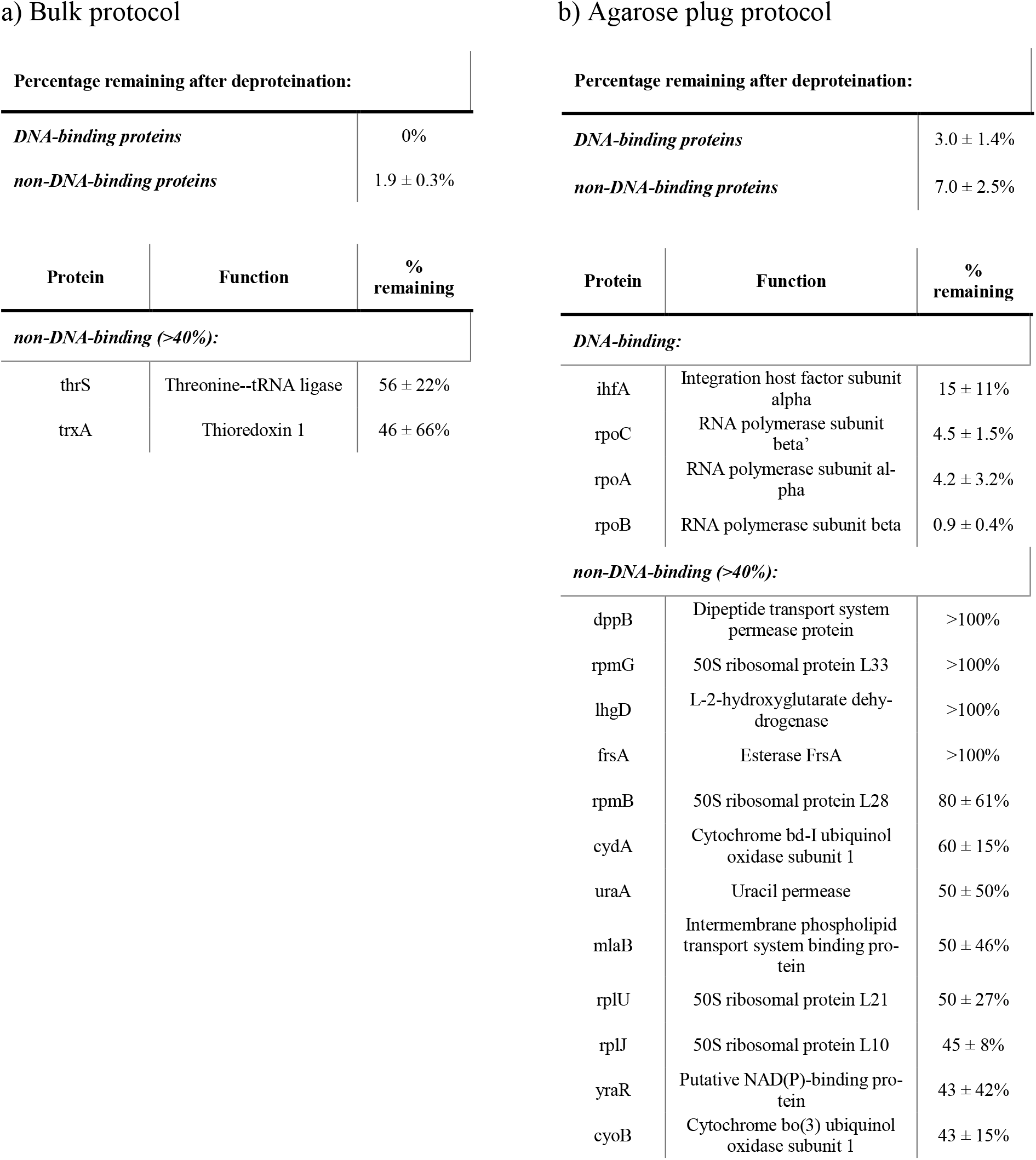
Protein removal efficiency as measured by mass spectrometry. **a)** Top: Percentage of proteins (total and DNA-binding proteins) remaining after the protein removal treatment for bulk protocol. Bottom: individual remaining proteins in the bulk protocol. *All* remaining DNA-binding protein are included, while for non-DNA-binding proteins only those with more than 40% remaining are included in the table. **b)** Same for the agarose plug protocol. The agarose plug protocol contained a few lower abundant proteins (dppB, rpmG, IhgD, frsA) for which higher relative abundancies were estimated (denoted with >100%) due to low level of protein removal. Errors are standard deviation from the mean obtained from three independent experiments per condition (‘before’ and ‘after’).

### Extracted chromosomes remain of megabasepair length and expand in size after protein removal

We imaged DNA resulting from the bulk and agarose plug protocols before and after protein removal by fluorescence imaging on a spinning disc confocal microscope using the DNA-intercalating dye Sytox-Orange (figure 3c/d and figure S2). From a first visual inspection we observed that, before protein removal, the DNA objects contain a dense/bright core with a lower density ‘cloud’ surrounding it (figure 3c-purple, figure 3d-orange/purple, and figure S2a/c/d). After protein removal, the objects seemed to be larger and more spread out (figure 3c/d-green, and figure S2b/e). In order to make more quantitative statements, we developed a semi-automated analysis script in Python (see Methods for a detailed description), with which we identified individual DNA objects in the images, segmented them from the background, and quantified their radius of gyration *R_g_* (a measure of the spatial extent of a polymer) as well as the sum of the fluorescence intensity.

**Figure 3.**
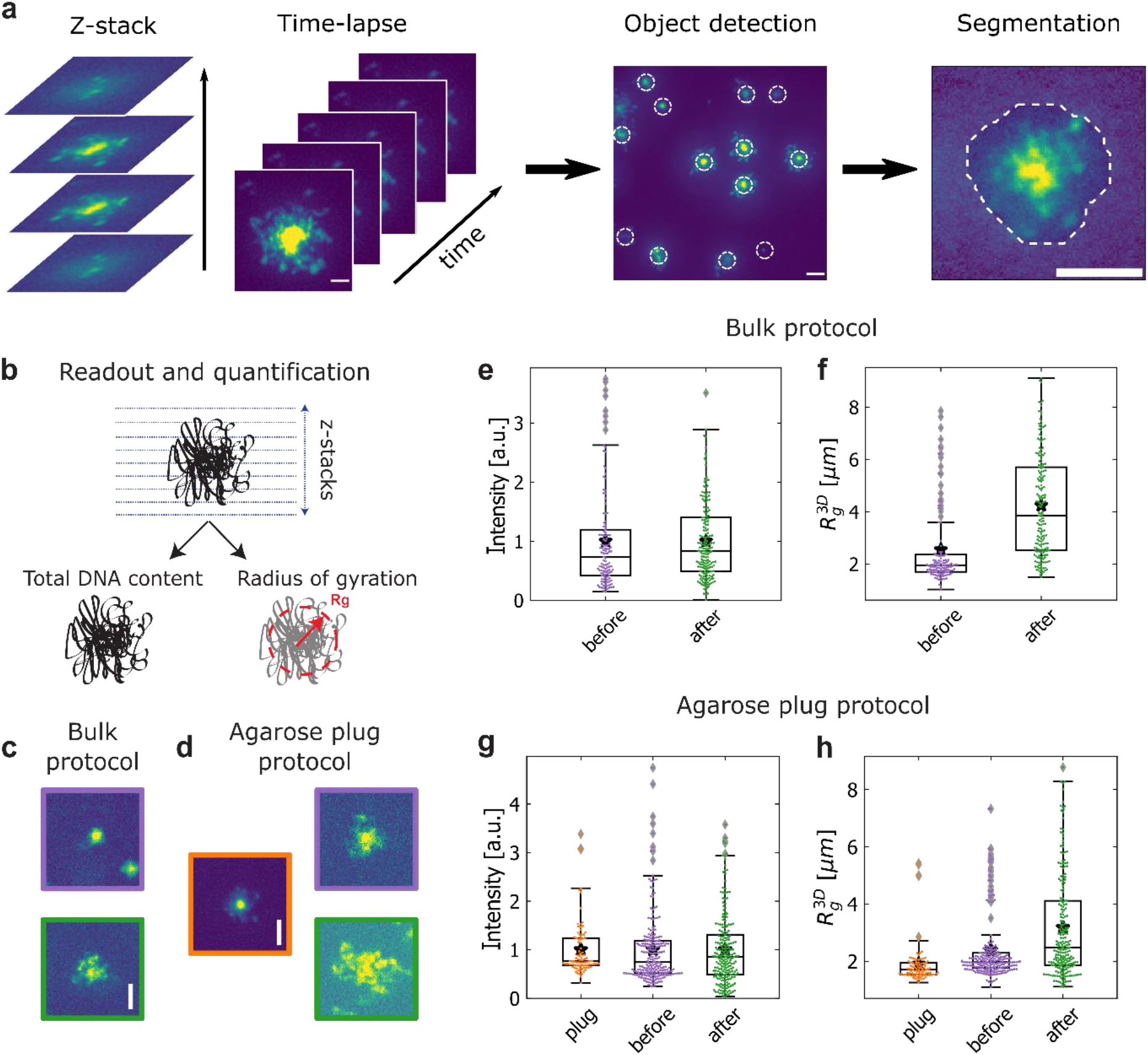
Characterization of isolated chromosomes before and after protein removal. **a)** Image analysis workflow for a GenBox experiment. In each image, objects are detected and segmented from the background. **b)** Within the segmentation boundary of each DNA object, the *R_g_* and the total fluorescence intensity are calculated. **c)** Images of typical DNA objects before (violet) and after (green) protein removal. **d)** Images of typical DNA objects in each condition of the agarose protocol: in plug (orange), before (violet) and after (green) protein removal. **e)** Total fluorescence intensity per DNA object before and after protein removal for the bulk protocol. **f)** *R_g_* distribution before and after protein removal for the bulk protocol. **g)** Total fluorescence intensity per DNA object before (in the plug and after plug melting) and after protein removal for the agarose plug protocol. **h)** *R_g_* distribution in the plug (orange), before protein removal but plug melting (purple) and after protein removal (green) for the agarose plug protocol. Boxplots show the median and 25^th^-75^th^ percentiles, star denotes mean. Scale bars are 5 μm. Intensity values in each distribution in **e** and **g** are scaled to the mean of the respective sum intensity distribution. Sample sizes in **e**, **f)** are N=125 and 181 for before and after. Sample sizes in **g**, **h)** are N=90, 223, 222 for plug, before, and after.

In our image analysis, the positions of DNA-objects were automatically determined from three-dimensional *z*-stacks followed by a manual curation step (figure 3a-object detection). Objects were then segmented in cube-shaped crops centered at each object’s center of mass. The DNA objects were further segmented from background within these cubes based on a globally (within the cube) determined threshold (Vtyurina, 2016), yielding a 3-dimensional foreground mask containing only the DNA object, and a minimal amount of background (figure 3a-segmentation and figure S4b). Masks determined on the individual crops were registered within the full field-of-view volume resulting in a labeled image. Individual masks were additionally checked in a curation step and manually adjusted if upon visual inspection they did not contain single objects or did not mask objects in their entirety. Sum intensity was calculated as the total sum of all pixel intensities within a foreground mask and the radius of gyration was calculated by squaring the sum of all foreground pixels’ intensity-weighted distances from the object’s center of mass (figure 3b) (Strychalski et al., 2012).

In order to monitor the integrity of the genomes at various steps of the protocol, we measured the total per-object fluorescence intensity, *i.e*. the sum of the intensities across all layers of the *z-*stack. While the sum intensity of a DNA object is expected to be set by the number of DNA basepairs, the measured distributions appeared to be fairly broad. In order to best compare the distributions before and after protein removal, we scaled the sum intensity values of each distribution with the mean value. We assume that the points in the ‘before’-distributions (before protein removal) in figure 3e and 3g represented those of intact chromosomes. This appears to be is a reasonable assumption since we observed similarly broad distributions of the sum intensity for lambda (λ)-DNA molecules (Figure S5).

To estimate the fraction of chromosomes that got fragmented in the process, we counted the objects in the distributions after protein removal that had a lower sum intensity value than a threshold of 1.5 times below the 25th percentile of the data. For the bulk protocol, this fraction was 4 of 181 objects, while for the agarose plug protocol it was 24 of 222 objects. In other words, only a low percentage of fragmented objects of 2% and 11% was estimated for bulk and agarose plug protocol, respectively. Another indication that our observed DNA objects remain well contained in the megabasepair size range comes from comparing their sum intensities with those of λ-DNA molecules (Table S3). We found that the mean of the ‘after’ sum intensity distribution is a factor 50 (bulk protocol) or 64 (agarose plug protocol) larger than the mean of the sum intensity distribution of the 48.5 kbp long lambda-DNA molecules. Assuming that the sum intensity scales linearly with the number of basepairs, this indicates that the DNA objects after protein removal have an average length of 2.4 Mbp (bulk protocol) and 3.1 Mbp (agarose plug protocol). However, these numbers are lower limits and the molecules are likely larger, because, following the same calculation, even the in-plug 4.6 Mbp chromosomes, which clearly are not fragmented, would be estimated to be 3.5 Mbp long.

The effect of deproteination of the genomes is also evident from an expansion in the size of the DNA objects, which can be characterized by measuring its radius of gyration. The mean *R_g_* in the bulk protocol increased from 2.55 ± 0.14 μm to 4.24 ± 0.14 μm (mean ± S.E.M) before and after protein removal respectively (Fig. 3f), and from 2.38 ± 0.08 μm to 3.18 ± 0.12 μm for the agarose plug protocol (figure 3h). These results indicate that the removal of the proteins had a clear effect on the mean *R_g_*, namely a 35% to 65% increase of the size for the agarose plug and bulk protocols, respectively. The measured radii of gyration exhibited a rather broad distribution (figure 3f/h). Notably, the measured *R_g_* values are extracted from momentarily measured snapshot images of the DNA objects, which yielded a broader distribution than the single value for the theoretical radius of gyration of a polymer which is a steady-state property (de Gennes, 1979).

### First proof-of-principle GenBox experiments

In order to demonstrate the potential of the GenBox approach, some first example experiments were performed. First, purified protein LacI was added to chromosomes that were deproteinated with the agarose plug protocol. These fluorescently labelled proteins bind sequence-specifically to FROS arrays that were inserted near the Ori position of the chromosomes. This yielded a well-visible fluorescent spot on the isolated chromosome (figure 4a-*ii*). Using a custom tracking script, the spot’s locations were tracked and the mean square displacement (MSD) was computed (figure 4a-*iii*). In line with the literature of local motion of chromosomal loci (Javer et al., 2014; Weber et al., 2010), the data for this example indicate that the DNA locus moved in a sub-diffusive manner, as the MSD curve tended to plateau towards longer lag times.

**Figure 4.**
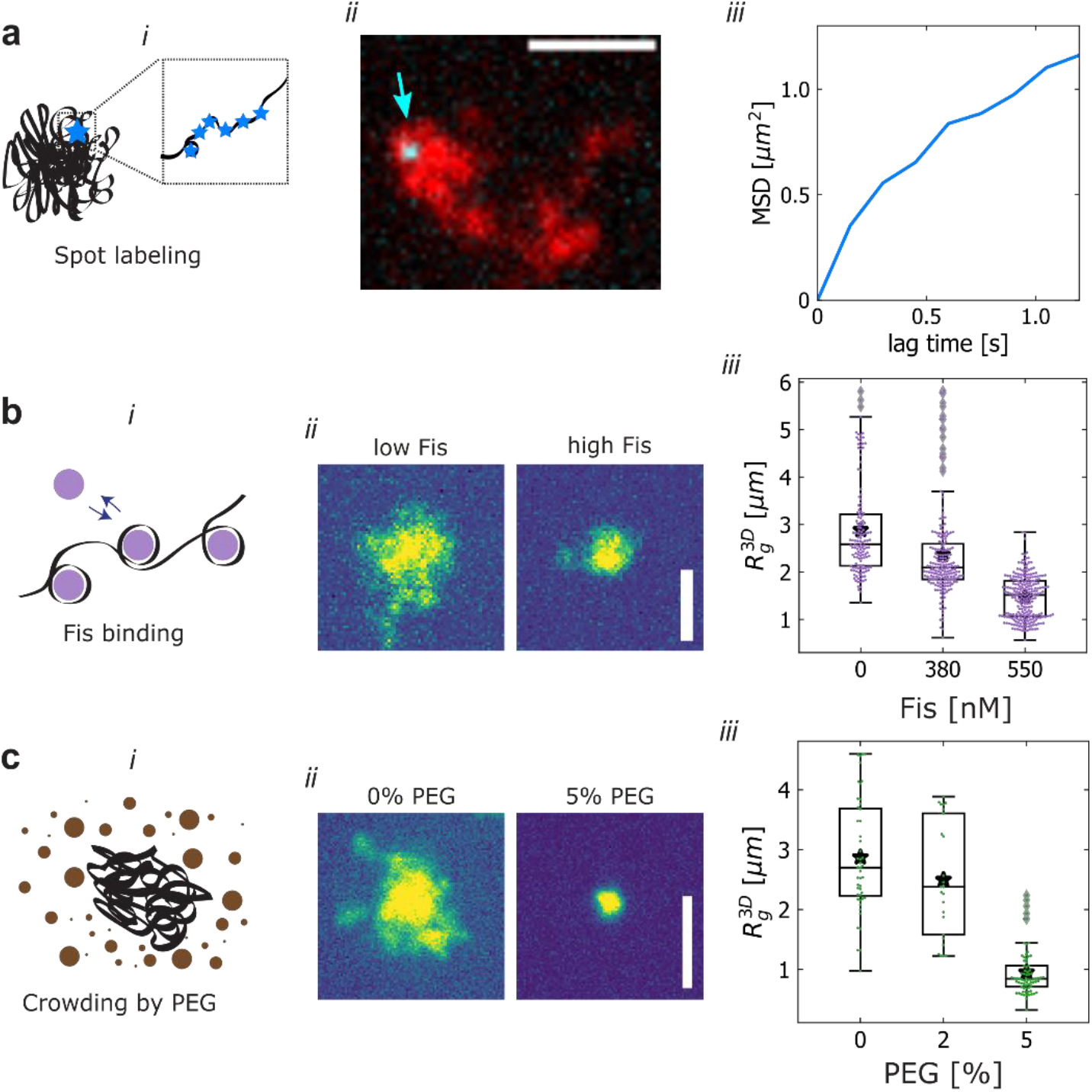
Proof-of-concept GenBox experiments. **a)** Example of a fluorescent spot located near the Ori (cyan). Location on the isolated chromosome (red) is tracked, yielding the MSD versus time (right) (Vink et al., 2020). **b)** Fis protein is added at increasing concentrations of 380 nM and 550 nM, and the resulting compaction is observed in the shifting and narrowing distribution of *R_g_* (right). **c)** PEG crowding agent is added at increasing concentrations of 2% and 5% and the resulting compaction is observed from the shifting and narrowing distribution of *R_g_*. Boxplots show the median and 25^th^-75^th^ percentiles, star denotes mean. Sample sizes are N=141, 201 and 242 in **b)** and N=48, 25, 74 in **c)**. All scale bars are 5 μm.

For a second example, the DNA-binding protein Fis was added to deproteinated chromosomes. Figure 4b-*ii* shows an example of a typical DNA object before and after addition of 550 nM Fis. A significant compaction of the DNA upon Fis addition is clear. The distributions of *R_g_* can be used to quantify the level of DNA compaction at increasing levels of added Fis (figure 4b-*iii*). As the Fis levels increased from 0 nM to 550 nM, the average *R_g_* decreased from 2.89 ± 0.08 μm to 1.47 ± 0.03 μm (mean ± S.E.M.), while the standard deviation of the distribution also decreased significantly from 1.00 μm to 0.45 μm.

For a final example, the crowding agent PEG was added at increasing concentrations to deproteinated chromosomes. A pronounced compaction was observed, when adding 5% PEG (figure 4c-*ii*), consistent with previous reports (Pelletier et al., 2012; Wegner et al., 2016). The increase of PEG from 0% to 2% resulted in the mean *R_g_* decreasing slightly from 2.87 ± 0.14 μm to 2.5 ± 0.2 μm, while the standard deviation remained steady at around 0.95 μm. However, at 5% PEG the mean and standard deviation of the *R_g_* distribution dropped to 0.95 ± 0.05 μm and 0.39 μm, respectively (figure 4c-*iii*).

## Discussion

In this paper, we present a methodology to prepare megabasepair deproteinated DNA, characterized the resulting DNA objects, and we provide first proof-of principle experiments to illustrate the utility of the method. The work expands on previous *in vitro* studies of large DNA molecules. For example, Wegner *et al*. (Wegner et al., 2012, 2016) and Cunha *et al*. (Cunha et al., 2001, 2005) studied bacterial chromosomes directly after isolation from cells in an aqueous solution, while Pelletier *et al*. (Pelletier et al., 2012) used microfluidic devices to perform cell lysis on-chip in cell-sized channels for studying the compaction of DNA with crowding agents. A limitation of these interesting first studies was that the megabasepair DNA substrates still contained an unknown number of natively bound proteins. Our GenBox protocol builds upon these previous experiments by explicitly removing the proteins and characterizing the remaining protein content with mass spectrometry and quantitative fluorescence imaging.

We presented two variants to prepare the deproteinated DNA sample, namely the bulk protocol and the agarose plug protocol. From a practical point of view, the agarose plug protocol has some advantages compared to the bulk protocol. First, samples can be made in advance and stored until needed for further processing. Secondly, unlike the bulk protocol sample, the agarose plugs are compatible with protocols that necessitate washing steps. One the other hand, the main advantage of the bulk protocol is the lower number of experimental steps. Our mass spectrometry data (table 1) showed that the deproteinated chromosomes of the bulk protocol contained fewer remaining DNA-binding proteins than those resulting from the agarose plug sample (0% vs 3%). Additionally, the bulk protocol results in a lower amount of fragmentation compared to the agarose plug protocol (as 98% *vs*. 89% of DNA objects classified as intact after protein removal). Since long DNA is easily sheared, it is important to limit the number of pipetting steps of DNA in solution. For both the bulk and agarose plug protocol, there is one major pipetting step involving the long DNA, namely the transfer to the observation well before the protein removal treatment. Conducting the chromosome extraction and protein removal inside a microfluidic chip could possibly eliminate this single pipetting step to further increase the number of intact DNA objects.

Modelling would be welcome to describe the observed radius of gyration of the deproteinated genomes. Polymer models connect the DNA contour length to a radius of gyration *R_g_* of the polymer blob that it forms in solution, but a broad spectrum of model variants that have been reported in literature yielded widely ranging values for *R_g_*. Indeed, how the theoretical *R_g_* scales with polymer length depends on multiple external parameters (de Gennes, 1979). These include, but are not limited to experimental parameters such as the fluorescent dyes (Japaridze et al., 2015), buffer salts and divalent cations, which set the solvent conditions and the resulting self-avoidance/attraction of the polymer, as well branches in the form of supercoils, the DNA topology of linear *vs* circular polymers, *etc*. Variation of these factors can yield very different predicted values for *R_g_* ranging from 1 to 6 μm for 4.6 Mbp DNA, as illustrated in Table S1. The values of *R_g_* that we observed in our experiments fall within this range. Notably, bacterial chromosomes may be natively supercoiled (Kavenoff and Bowen, 1976). While the removal of supercoil-stabilizing proteins as well as potential local nicks in the DNA will likely reduce the level of supercoiling significantly, some degree of supercoiling may remain in the DNA objects that result from the protocol.

We hope that the results presented in this paper open a way to start GenBox experiments that may subsequently provide a valuable bottom-up approach to the field of chromosome organization. Promising avenues may include encapsulation of megabasepair DNA inside droplets or liposomes to study the effects of spatial confinement, addition of loop extruding proteins such as cohesin or condensin to elucidate the effect of loop formation on the structure of large DNA substrates, and experiments with phase-separating DNA-binding proteins to observe the effects of polymer-mediated phase separation at long length scales.

## Materials and Methods

### Preparation of spheroplasts and imaging of cells and ori/ter ratio

*E. coli* bacterial cells (HupA-mYPet frt, Ori1::1acOx240 frt, ter3::tetOx240 gmR, ΔgalK::tetR-mCerulean frt, ΔleuB::lacI-mCherry frt, DnaC::mdoB::kanR frt)(Wu et al., 2019) were incubated from glycerol stock in M9 minimal media (1x M9 minimal salts, 0.01 % v/v protein hydrolysate amicase, 0.8% glycerol, 0.1 mM CaCl_2_, 2 mM MgSO_4_) supplemented with 50 μg/mL Kanamycin antibiotic (K1876, Sigma-Aldrich) in a shaking incubator at 30 °C and 300 rpm and allowed to reach OD_600_ of 0.1 to 0.15. The cells were then grown for 2 to 2.5 hours at 41 °C shaking at 900 rpm in order to arrest replication initiation.

In order to determine the Ori/Ter ratio, 1.25 μL cells were deposited on a cover slip (15707592, Thermo Fischer) and covered with an agarose pad. The cells were imaged with a Nikon Ti2-E microscope with a 100X CFI Plan Apo Lambda Oil objective with an NA of 1.45 and SpectraX LED (Lumencor) illumination system using phase contrast, cyan (CFP filter cube λ_ex_/λ_bs_/λ_em_ = 426–446/455/460–500 nm), yellow (triple bandpass filter λ_em_ = 465/25–545/30–630/60 nm) and red (the same triple bandpass filter). Spots corresponding to Ori and Ter were identified on the red and cyan channels and counted either manually or with an automated routine, producing the same results.

Next, appropriate volume of cell culture was spun down at 10000 g for 2.5 min, in order to obtain a pellet at OD_eq_ = 1 (approx. 8 x 10^8^ cells). The pellet was resuspended in 475 μL cold (4 °C) sucrose buffer (0.58 M sucrose, 10 mM Sodium Phosphate pH 7.2, 10 mM NaCl, 100 mM NaCl). 25 μL lysozyme (L6876 Sigma-Aldrich, 1 mg/mL in ultrapure water) was immediately added and gently mixed into the cell/sucrose buffer suspension, followed by either *i*) 15 min incubation at room temperature (bulk protocol) or *ii*) *a* 10 min incubation at room temperature and a 5 min incubation at 42 °C in a heat block (agarose plug protocol). The lysozyme digests the cell wall, resulting in spheroplasts.

### Preparation of isolated chromosomes (bulk protocol)

Spheroplasts were prepared as described above. Cell lysis and nucleoid release was achieved by pipetting 10 μL of spheroplasts into 1 mL of lysis buffer (20 mM Tris-HCl pH 8) with a cut pipette tip, after which the tube was once gently inverted for mixing. Immediately thereafter, buffer composition was adjusted to match the one of agarose plug protocol (50 mM Tris-HCl pH 8, 50 mM NaCl, 1 mM EDTA pH 8.0 and 5% glycerol). After this stage, we continued to the preparation of the observation well.

### Preparation of isolated chromosomes (agarose plug protocol)

500 μL warmed (42 °C) spheroplast/sucrose buffer suspension was added to 500 μL warm (42 °C) agarose solution (low melting point agarose, V2831 Promega, 2% w/v in sucrose buffer) using a cut pipette tip. In the following steps, the Eppendorf tubes were kept at 42 °C to prevent gelation of the agarose solution. The spheroplast/agarose mixture was gently mixed using a cut pipette tip, and casted in volumes of 100 μL into a plug mold (Bio-Rad laboratories, Veenendaal, The Netherlands). In order to produce a larger number of agarose plugs, it proved most optimal to perform the protocol with multiple Eppendorf tubes in parallel, rather than increasing the number of cells and volumes of sucrose buffer and agarose solution used per Eppendorf tube. To solidify the agarose plugs, the plug mold was stored at 4 °C for 1 h.

The solidified agarose plugs containing spheroplasts were removed from the plug mold and added to 25 mL per plug lysis buffer (10 mM Sodium Phosphate pH 7.2, 10 mM EDTA pH 8.0, 100 μg/mL RNase-A), thereby lysing the cells and thus merely trapping the nucleoids from the spheroplasts in the agarose gel matrix. The plugs were incubated tumbling in the lysis buffer for 1 h. Subsequently, the plugs were removed from the lysis buffer and each plug was stored in 2 mL TE wash buffer (20 mM Tris-HCl pH 8, 50 mM EDTA pH 8.0) at 4 °C until further use.

In order to transfer agarose plugs from one container to another, a sheet of aluminum foil was put over the top of a glass beaker. Using a 200 μL pipette tip holes were punched into the aluminum foil and the foil was gently pressed down into a concave shape to prevent liquid spilling over the edge. The container containing the plugs was emptied through the strainer into the beaker, leaving the agarose plugs behind on the strainer. Using flat-headed tweezers the agarose plugs were transferred to the new container. To prevent cross-contamination, the tweezers were washed after each handing step with 70% ethanol and dried using a pressurized air gun.

For releasing the purified genome from the agarose plugs for experiments, agarose plugs were incubated for 1 hour in buffer A (50 mM Tris-HC pH 8, 50 mM NaCl, 1 mM EDTA pH 8.0, 5% glycerol) and then transferred to 150 μL of buffer A preheated to 71 °C. The plug was then melted at 71 °C for 15 minutes before equilibrating at 42 °C. The agarose was digested by 1 hour incubation at 42 °C with 2 units of beta-agarase (M0392, New England Biolabs). After this stage, we continued to the preparation of the observation well.

### Imaging of spheroplasts and chromosomes inside the agarose plug

A plug containing spheroplasts was deposited on a KOH-cleaned cover slip. Spheroplasts were imaged with a Nikon Ti2-E microscope with a 100X CFI Plan Apo Lambda Oil objective with an NA of 1.45 and SpectraX LED (Lumencor) illumination system using the channels phase contrast, cyan (CFP filter cube λex/λbs/λem = 426–446/455/460–500 nm), yellow (triple bandpass filter λem = 465/25–545/30–630/60 nm) and red (the same triple bandpass filter). The imaging protocol was composed of a single time-point, using a 2 μm z-stack with 200 nm z-slices.

For imaging chromosomes after lysing the spheroplasts, a nucleoid-containing plug was incubated in 2 mL buffer A (50 mM Tris-HC pH 8, 50 mM NaCl, 1 mM EDTA pH 8.0, 5% glycerol) at 4 °C for 1 h. The plug was transferred to 2 mL imaging buffer (50 mM Tris-HC pH 8, 50 mM NaCl, 1 mM EDTA pH 8.0, 5% glycerol, 3.5 mM MgCl_2_, 1 mM DTT, 500 nM Sytox Orange) and incubated for 15 min. Then the plug was deposited on a KOH-cleaned cover slip and 30 μL imaging buffer was added onto the plug to prevent drying. The plug was imaged using an Andor Spinning Disk Confocal microscope with a 100x oil immersion objective, 20% 561 laser, filters, 250x gain, and 10 ms exposure. The imaging protocol resulted in 30 μm z-stacks with 250 nm z-slices and was repeated at 15 distinct XY positions.

### Treatment with Proteinase K for protein removal

Thermolabile Proteinase K (P8111S, New England Biolabs) was added to isolated chromosomes (0.01 unit per 1 μL of nucleoid suspension) in buffer containing 2.5 mM MgCl_2_ and 50 mM NaCl. The samples were then incubated for 15 minutes at 37 °C for treatment and for 10 minutes at 56 °C for Proteinase K inactivation. The samples were equilibrated to RT for at least 30 minutes before imaging or and further experiments.

### Mass spectrometry

Bulk and agarose plug samples were treated with Proteinase K as described above. Each sample contained nucleoids from an amount of cells corresponding to OD 5.0 (ca. 5×10^9^ cells in 100 μL). With two different DNA isolation approaches (bulk and agarose plug) and two conditions (control and Proteinase K) four triplicate samples were analyzed (twelve samples in total) by mass spectrometry. Control sample was underwent exactly the same steps as the treated sample, but equal volume of 50 % glycerol (corresponding to Proteinase K storage buffer concentration) was used instead of Proteinase K enzyme. 200 mM ammonium bicarbonate buffer (ABC) was prepared by dissolving ammonium bicarbonate powder (A6141, Sigma-Aldrich) in LC-MS grade quality water. 10 mM DTT (43815, Sigma-Aldrich) and iodoacetamide (IAA) (I1149, Sigma-Aldrich) solutions were made fresh by dissolving stock powders in 200 mM ABC. Next, 25 μL of 200 mM ABC buffer was added to each sample to adjust pH, immediately followed by addition of 30 μL of 10mM DTT and 1 hour incubation at 37 °C and 300 rpm. Next, 30 μL of 20 mM IAA was added and samples were incubated in dark at room temperature for 30 min. Finally, 10 μL of 0.1 mg/mL trypsin (V5111, Promega) was added and samples were incubated overnight at 37 °C and 300 rpm.

On the following day, samples were purified by solid phase extraction (SPE). SPE cartridges (Oasis HLB 96-well μElution plate, Waters, Milford, USA) were washed with 700 μL of 100% methanol and equilibrated with 2×500 μL LC-MS grade H_2_O. Next, 200 μL of each sample was loaded to separate SPE cartridge wells and wells were washed sequentially with 700 μL 0.1% formic acid, 500 μL of 200 mM ABC buffer and 700 μL of 5% methanol. Samples were then eluted with 200 μL 2% formic acid in 80 % methanol and 200 μL 80% 10 mM ABC in methanol. Finally, each sample was collected to separate low-binding 1.5 μL tubes and speedvac dried for 1-2 hours at 55 °C. Samples were stored frozen at −20 °C until further analysis. Desalted peptides were reconstituted in 15 μL of 3% acetonitrile/0.01% trifluoroacetic acid prior to MS-analysis.

Per sample, 3 μL of protein digest was analysed using a one-dimensional shotgun proteomics approach (Köcher et al., 2012; den Ridder et al., 2022). Briefly, samples were analysed using a nano-liquid-chromatography system consisting of an EASY nano LC 1200, equipped with an Acclaim PepMap RSLC RP C18 separation column (50 μm x 150 mm, 2 μm, Cat. No. 164568), and a QE plus Orbitrap mass spectrometer (Thermo Fisher Scientific, Germany). The flow rate was maintained at 350 nL·min^-1^ over a linear gradient from 5% to 25% solvent B over 90 min, then from 25% to 55% over 60 min, followed by back equilibration to starting conditions. Data were acquired from 5 to 175 min. Solvent A was H_2_O containing 0.1% FA, and solvent B consisted of 80% ACN in H_2_O and 0.1% FA. The Orbitrap was operated in data-dependent acquisition (DDA) mode acquiring peptide signals from 385–1250 m/z at 70,000 resolution in full MS mode with a maximum ion injection time (IT) of 75 ms and an automatic gain control (AGC) target of 3E6. The top 10 precursors were selected for MS/MS analysis and subjected to fragmentation using higher-energy collisional dissociation (HCD). MS/MS scans were acquired at 17,500 resolution with AGC target of 2E5 and IT of 100 ms, 1.0 m/z isolation width and normalized collision energy (NCE) of 28.

Data were analysed against the proteome database from *Escherichia coli* (UniProt, strain K12, Tax ID: 83333, November 2021, https://www.uniprot.org/), including Proteinase K from *Parengyodontium album* (UniProt ID: P06873) and Beta-agarase I from *Pseudoalteromonas atlantica* (UniProt ID: Q59078) (Bateman et al., 2017), using PEAKS Studio X+ (Bioinformatics Solutions Inc., Waterloo, Canada) (Ma et al., 2003), allowing for 20 ppm parent ion and 0.02 m/z fragment ion mass error, 3 missed cleavages, carbamidomethylation as fixed and methionine oxidation and N/Q deamidation as variable modifications. Peptide spectrum matches were filtered for 1% false discovery rates (FDR) and identifications with ≥ 1 unique peptide matches. For the case that a protein in the sample was identified by only a single peptide in only one out of three runs, the protein identification was only considered if the same peptide sequence was also identified in unpurified control (within a retention time window of ± 2 min). For determination of relative amounts of protein remaining after Proteinase K treatment, protein abundances were expressed as ‘spectral counts’ normalized by their molecular weight (*i.e*., 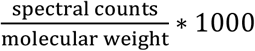). Using the normalized spectral counts per protein in the three replicate experiments per condition (‘before’ and ‘after’), the mean was calculated for each protein individually and for the aggregated DNA-binding and non-DNA-binding categories. Uncertainties were expressed as standard deviations from the means due to inter-sample variation. Relative amounts (for individual proteins and the aggregated categories) were defined as the ratio of the ‘after’ over the ‘before’ means, with uncertainties calculated by propagating the errors through this ratio.

### Preparation of observation wells

Cover slips (15707592, Thermo Fischer) were loaded onto a teflon slide holder. The coverslips were sonicated in a bath sonicator in a beaker containing ultrapure water for 5 min, followed by sonication in acetone for 20 min, a rinse with ultrapure water, sonication in KOH (1 M) for 15 min, a rinse with ultrapure water, and finally sonication in methanol for 15 min. Cleaned cover slips were stored in methanol at 4 °C.

To assemble the observation well, a PDMS block with a 4 mm punched (504651 World Precision Instruments) through hole was bonded on a cleaned coverslip. PDMS block was obtained from PDMS slab of ± 5 mm thickness which was casted from mixture of 10:1 = PDMS:curing agent (Sylgard 184 Dow Corning GmbH) and allowed to cure for 4 hours at 80 °C. The bonding was done immediately after exposing both surfaces, glass and PDMS, to oxygen plasma (2 minutes at 20 W) and the bond was allowed to cure for 10 minutes at 80 °C.

Immediately after the bonding, the inner surface of the observation well was treated to create a lipid bilayer to prevent sticking of DNA and proteins. To do so, DOPC liposomes were used. DOPC and PE-CF lipids from chloroform stocks (both Avanti Polar Lipids, Inc.) were combined in 999:1 mol-ratio DOPC:PE-CF in a glass vial for final lipid concentration of 4 mg/mL. Chloroform was evaporated by slowly turning the vial in a gentle nitrogen steam for 15 minutes or until dry. The vial was then placed in a desiccator for 1 hour to further dry its contents. The lipids were then resuspended in SUV buffer (25 mM Tris-HCl pH 7.5, 150 mM KCl, 5 mM MgCl_2_) and vortexed until solution appears opaque and homogeneous to the eye. Any large lipid aggregates were broken up by 7 to 10 freeze-thaw cycles of repeated immersion into liquid nitrogen and water at 70-90 °C. The lipid suspension was loaded in a glass syringe (250 μL, Hamilton) and extruded through 30 nm polycarbonate membrane (610002, Avanti Polar Lipids, Inc.) fixed in mini-extruder (610020, Avanti Polar Lipids, Inc.) at 40 °C. Lipids were stored at −20 °C for up to several months. SUV suspension (99.9 mol% DOPC, 0.1 mol% PE:CF - both Avanti Polar Lipids, Inc.) was sonicated for 10 minutes at RT and pipetted into the well to cover the area to be treated. After 1 minute of incubation, the solution was diluted by adding 3x fold excess off SUV buffer (25 mM Tris-HCl pH 7.5, 150 mM KCl, 5 mM MgCl_2_). Subsequently, the solution in the well was exchange at least 5-times, without de-wetting the surface of the glass, for imaging buffer (50 mM Tris-HC pH 8, 50 mM NaCl, 1 mM EDTA pH 8.0, 5% glycerol, 3.75 mM MgCl_2_, 1.5 mM DTT, 750 nM Sytox Orange). As final step, a sample with nucleoids from either the bulk or plug protocol was added to the imaging buffer in ratio 1:2 (nucleoids to imaging buffer), after which the well was ready for imaging.

### Experiments with spot labeling, Fis, and PEG

For the experiments of Fig.5, the protocol for imaging digested plugs was followed, but with some modifications for the imaging. Plugs with ProtK protein removal treatment were used. The imaging protocol was as follows: *i*) a 30 μm z-stack was taken with 250 nm z-slices, and this was repeated at 5 XY positions; *ii*) a 30 μm z-stack was taken with 1 μm z-slices at 5 XY positions, repeated 10 times; *iii*) the protein of interest was added to the observation well at a final concentration of 1.25 nM (LacI), 380 or 550 nM (Fis), 2 or 5% (PEG-8000, Sigma Aldrich); *iv*) a 30 μm z-stack was taken with 1 μm z-slices at 5 XY positions, repeated 50 times. Once the compaction process reached a steady state, the imaging step *i*) was repeated.

Fis protein was a kind gift of William Nasser, and was purified as described previously (Japaridze et al., 2021). 8xHis-tagged LacI-SNAP fusions in pBAD plasmids were ordered from GenScript. BL21(DE3)-competent *E.coli* cells (New England Biolabs) were transformed with the plasmids and plated with Ampicillin (Amp). Overnight colonies were inoculated in LB with Amp and incubated overnight at 37 °C and 150 rpm. Cells were diluted 1:100 into fresh media with Amp and grown at 37 °C at 150 rpm until OD_600_ of 0.5 - 0.6 after which 2 g/L arabinose was added to induce expression for 3-4 hours. Next, cells were harvested by centrifugation and resuspended in buffer A (50 mM Tris-HCl pH 7.5, 200 mM NaCl, 5 % w/v glycerol). Lysis was performed with French Press and supernatant was recovered after centrifugation. His-tagged proteins were bound to beads in talon resin and column was then in turns washed with 50 mL of buffer A1 (buffer A + 10 mM imidazole), buffer A2 (buffer A + 0.01 % Tween20), and buffer A3 (buffer A + 0.5 M NaCl). Next, the sample was eluted with 15 mL buffer B (buffer A + 3C protease + 1 mM β-Mercaptoethanol) and diluted 10x in buffer C (50 mM Tris-HCl pH 8.0). Anion exchange chromatography was done with Mono Q-ion exchange column (Cytiva) equilibrated with buffer C and sample was eluted to buffer D (50 mM Tris-HCl pH 8.0 with 1 M NaCl). Next, size exclusion chromatography was done on Superdex S200 (Cytiva) column equilibrated with buffer A, collected and fractions were run on gel to check for purity. Finally, purified proteins were labelled with SNAP-Surface Alexa Fluor 647 tag (New England Biolabs) following manufacturer’s instructions.

### Image processing and analysis

We developed a custom analysis pipeline for quantifying DNA objects in fluorescent images obtained from GenBox experiments, written entirely in Python. The analysis proceeds in three main steps: *i*) identification of individual DNA objects, *ii*) segmentation of these objects from background, *iii*) quantification of relevant observables (*e.g*., a calculation of the radius of gyration).

Positions of individual objects were determined automatically from threedimensional stacks using *skimage* function *peak_local_max* (Van Der Walt et al., 2014). Maxima were required to be at least twice as bright as globally determined threshold (Vtyurina, 2016) (see next paragraph for description). If objects’ maxima were closer than 30 pixels from each other, or from any image boundary, the objects were discarded from further analysis. Next, all locations were visually inspected with *napar*’s viewer (Sofroniew et al., 2022) using Image and Points layers. Typically, none or few changes had to be made (*e.g*., if one object was identified as two or vice-versa).

Objects were segmented from the background in crops corresponding to 25×25×25 μm^3^ centered at each object’s center of mass. First, the raw data in any crop was binarized based on a globally determined threshold (Vtyurina, 2016). Pixels’ intensity values were sorted increasingly, and two lines were fitted to such curve *a*) a line fitted to the first half of the pixels in the image (estimate of background), and *b*) a line fitted to all pixels brighter than half of the maximum intensity (estimate of foreground). The intensity threshold value was then determined from the point on the sorted intensity curve which was closest to intersection of the two lines (Fig. SI4a). Images before and after background subtraction were inspected and confirmed that the approach was able to discriminate background and foreground well (Fig. SI4b). The crops were then traversed plane-by-plane in *z-*direction, discarding small regions, dilating remaining region(s) and filling holes. The mask contours were smoothed in each plane with a Savitzky-Golay filter with a window size of quarter the contour length of the mask. Finally, only the most central 3D contiguous binary object was retained as foreground mask for each object.

Masks determined on individual crops were subsequently registered within full FOV volume (typically about 100×100×100 μm^3^) producing a labeled image. If shared pixels resulted at masks overlap, these pixels were assigned to the mask which center of mass was the closest. Subsequently, the masks were inspected with *napari’s* viewer using Image and Label layers and manually adjusted if upon visual inspection they did not contain single objects or did not mask those in their entirety.

The quantification of the objects’ properties was done within the volume of the foreground mask applied onto the raw data after subtracting globally determined threshold (as described earlier) from each crop. Sum intensity was calculated as the total sum of all pixel intensities within a foreground mask and the radius of gyration was calculated by squaring the sum of all foreground pixels’ intensity-weighted distances from the object’s center of mass. The resulting measurements were saved as structured JSON files, one per each FOV, and aggregated based on condition to produce R_g_ and intensity plots. The MSD in spot-labeling experiment was calculated using the *xy*-coordinates of fluorescent spots obtained with the ImageJ TrackMate plugin (Schindelin et al., 2012; Tinevez et al., 2017).

## Notes

### Competing Interest Statement

The authors have declared no competing interest.

