## Supplemental Tables and Figures for "Extracting and characterizing protein-free megabasepair DNA for *in vitro* experiments"

**Supplementary figures**

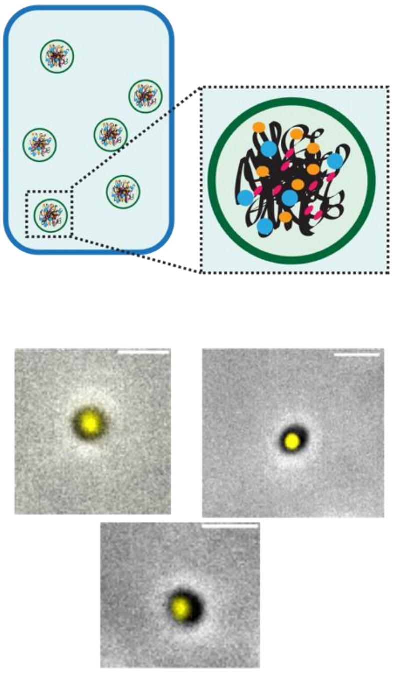

**Figure S1. Spheroplasts in plug**

Schematic (top) and microscopy images (bottom) of spheroplasts embedded inside an agarose plug. The yellow signal comes from fluorescently labeled HU-protein and thus serves as a DNA marker. The greyscale signal is phase contrast. Scale bars are 2 $\mu$m.

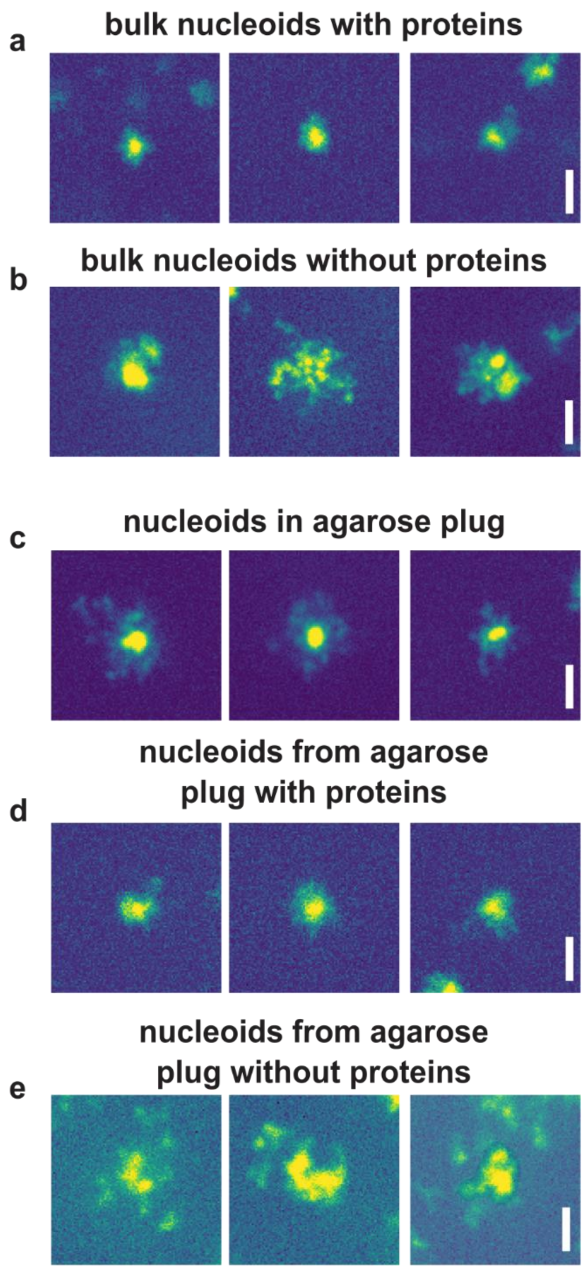


**Figure S2. Examples of DNA-objects**

Fluorescence images of DNA objects in various conditions: **a)** Bulk protocol chromosomes before protein removal. **b)** Bulk protocol chromosomes after protein removal. **c)** Agarose plug protocol chromosomes inside the agarose plug before protein removal. **d)** Agarose plug protocol chromosome in solution before protein removal. **e)** Agarose plug protocol chromosomes in solution after protein removal. Scale bars are 5 $\mu$m.

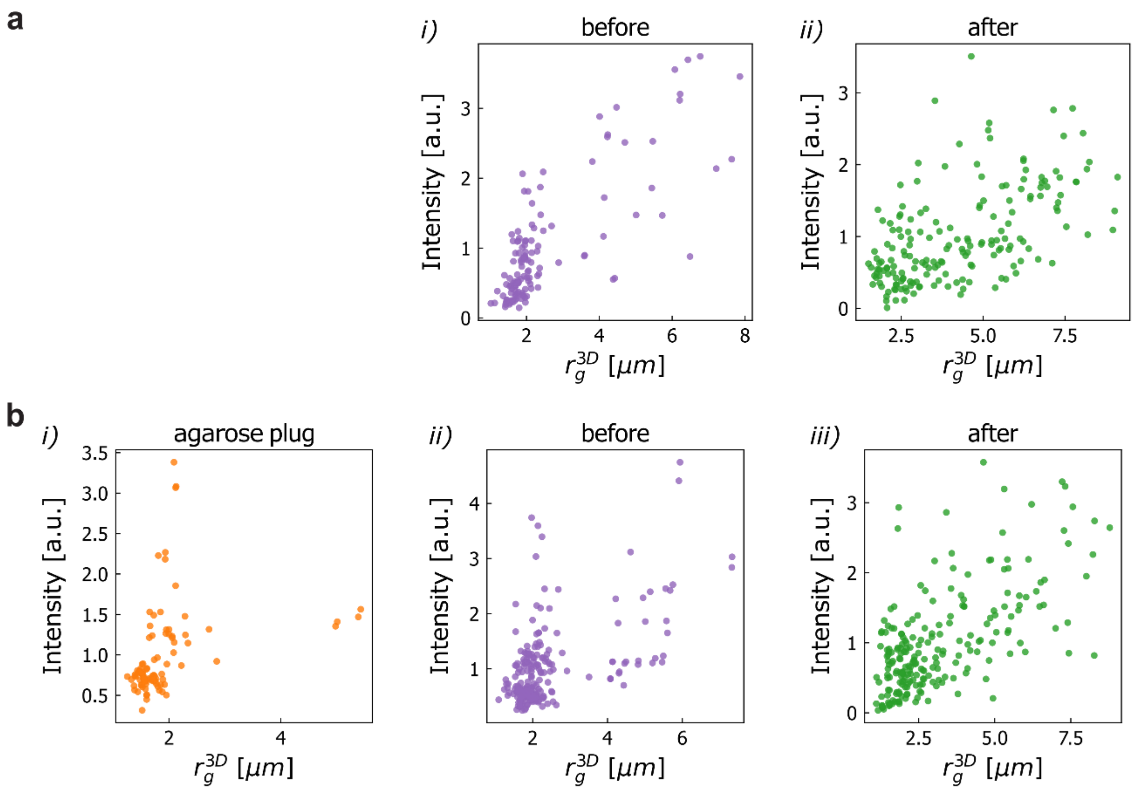


**Figure S3. Radius of gyration versus sum intensity distributions**

Scatter plots of the radius of gyration and sum intensity of observed DNA objects in various conditions: **a)** *i)* Bulk protocol chromosomes before protein removal. *ii)* Bulk protocol chromosomes after protein removal. **b)** *i)* Agarose plug protocol chromosomes inside the agarose plug before protein removal. *ii)* Plug protocol chromosome in solution before protein removal. *iii)* Agarose plug protocol chromosomes in solution after protein removal. Intensity values in each scatter plot are scaled to the mean of the applicable sum intensity distribution. Sample sizes are N=125 and 181 in **a)** and N=90, 223, 222 in **b)**.

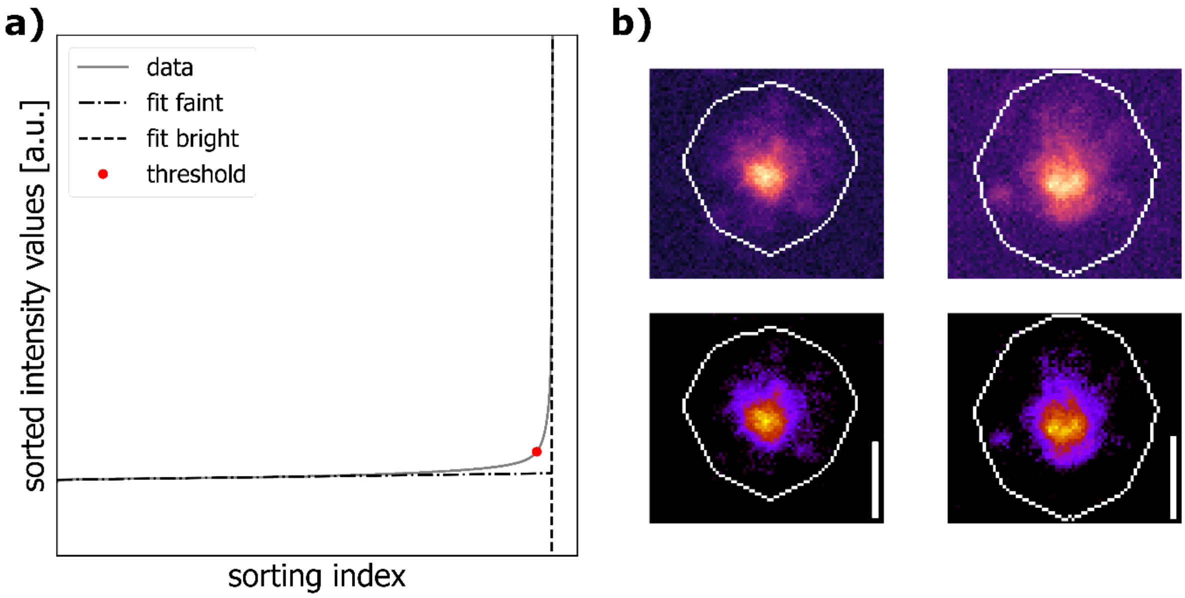


**Figure S4. Visualization of thresholding procedure**

**a)** Pixel intensity values were sorted by increasing intensity, and two lines were fitted to this curve: a line fitted to the first half of the pixels in the image (which is the estimate of background, dash-dot), and a line fitted to all pixels brighter than half of the maximum intensity (estimate of foreground, dash). The intensity threshold value was then determined from the point on the sorted intensity curve (red dot) which was closest to intersection of the two lines. **b)** Images before (top) and after (bottom) background subtraction. Inspection confirmed that the approach was able to discriminate background and foreground well. White line is contour of the mask. Scale bars are 5 $\mu$m.

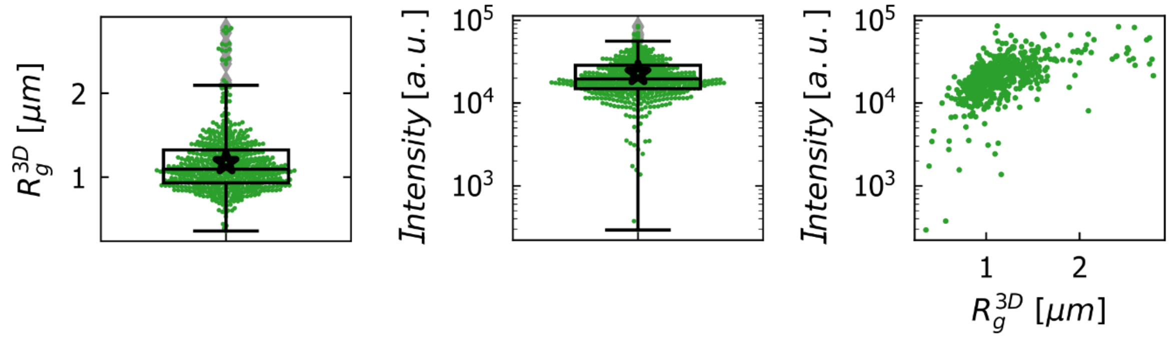


**Figure S5. Characterization of lambda-DNA molecules**

(*left*) $R_g$ distribution for lambda DNA molecules. (*center*) Total fluorescence intensity per identified λ-DNA molecule. (*right*) $R_g$ vs total fluorescence intensity per DNA object distribution. Boxplots show the median and 25th-75th percentiles, star denotes mean, N=534.

| | Morphology | Solvent | *N = 1 Mbp, $L_p$ = 25 nm [μm]* | *N = 1 Mbp, $L_p$ = 50 nm [μm]* | *N=4.6 Mbp, $L_p$ = 25 nm [μm]* | *N=4.6 Mbp, $L_p$ = 50 nm [μm]* |
|---|---|---|---|---|---|---|
| **Ideal chain** | L | n.a. | 1.7 | 2.4 | 3.6 | 5.1 |
| | R | n.a. | 1.2 | 1.7 | 2.6 | 3.6 |
| **Worm-like chain** | L | n.a. | 1.7 | 2.4 | 3.6 | 5.1 |
| | R | n.a. | 0.8 | 1.2 | 1.8 | 2.6 |
| **Self-avoiding polymer with solvent interaction (Flory theory)** | | good | 2.6 | 3.4 | 6.3 | 8.4 |
| | R | ideal | 1.2 | 1.7 | 2.6 | 3.6 |
| | | poor | 0.35 | 0.54 | 0.6 | 0.9 |
| | | good | 3.7 | 4.9 | 9.0 | 12.0 |
| | L | ideal | 1.7 | 2.4 | 3.6 | 5.1 |
| | | poor | 0.5 | 0.76 | 0.8 | 1.3 |
| **Uncrosslinked supercoiled polymer** | L/C | n.a. | 1.35 | 0.83 | 1.5 | 2.5 |

**Table S1. Gyration radii for various length DNA and various persistence length values.** The persistence length of bare DNA is commonly 50 nm. However due to the buffer conditions (salts, divalent cations, intercalation dyes) it can decrease to values as low as 25 nm. Morphology: L - linear, R - ring. Solvent: good - $\nu = 0.588$, ideal - $\nu = 0.5$, poor - $\nu = 0.36$.

| **Protein** | **Function** |
|---|---|
| rpoC | DNA-directed RNA polymerase subunit beta' |
| rpoB | DNA-directed RNA polymerase subunit beta |
| rpoA | DNA-directed RNA polymerase subunit alpha |
| gyrA | DNA gyrase subunit A |
| topA | DNA topoisomerase 1 |
| gyrB | DNA gyrase subunit B |
| stpA | DNA-binding protein StpA |
| hupA | DNA-binding protein HU-alpha |
| dps | DNA protection during starvation protein |
| ybiB | Uncharacterized protein |
| fis | DNA-binding protein Fis |
| cbpA | Curved DNA-binding protein |
| rpoZ | DNA-directed RNA polymerase subunit omega |
| polA | DNA polymerase I |
| hupB | DNA-binding protein HU-beta |
| ihfA | Integration host factor subunit alpha |
| ihfB | Integration host factor subunit beta |
| helD | DNA helicase IV |
| kdgR | Transcriptional regulator |
| uvrD | DNA helicase II |
| oxyR | Hydrogen peroxide-inducible genes activator |
| parE | DNA topoisomerase 4 subunit B |
| rpoS | RNA polymerase sigma factor |
| rpoD | RNA polymerase sigma factor |
| crl | Sigma factor-binding protein |
| yejK | Nucleoid-associated protein |
| ybaB | Nucleoid-associated protein |
| dnaE | DNA polymerase III subunit alpha |
| dnaA | Chromosomal replication initiator protein |
| ebfC | Nucleoid-associated protein |
| slmA | Nucleoid occlusion factor |
| crfC | Clamp-binding protein CrfC |
| mukB | Chromosome partition protein |
| mukF | Chromosome partition protein |
| matP | Macrodomain Ter protein |
| topo3 | DNA topoisomerase |
| parC | DNA topoisomerase 4 subunit A |
| mukE | Chromosome partition protein |

**Table S2. List of DNA-binding proteins used for mass spectrometry analysis**

Proteins’ description is taken UniProt (UniProt, strain K12, Tax ID: 83333, November 2021) database. Shortlist contains proteins identified as DNA-binding or DNA processing.

| sample | condition | intensity [a.u.] | relative intensity | (theoretical) number of kbp |
|---|---|---|---|---|
| **bulk** | after | 1'133'034 | 49.5 | 2'403 |
| **plug** | after | 1'466'184 | 64.1 | 3'109 |
| **plug** | in plug | 1'658'675 | 72.5 | 3'518 |
| **lambda** | | 22'870 | 1 | 48.5 |

**Table S3. Total and relative total intensities of DNA molecules**

Mean sum intensity per molecule is reported. Bulk and plug condition values are compared relative to lambda-DNA molecules. Shaded cell values are given by definition.
